# CellexalVR: A virtual reality platform to visualise and analyse single-cell data

**DOI:** 10.1101/329102

**Authors:** Oscar Legetth, Johan Rodhe, Stefan Lang, Parashar Dhapola, Joel Pålsson, Mattias Wallergård, Shamit Soneji

## Abstract

Single-cell RNAseq is a routinely used technique to explore the composition of cell populations, and they are often visualised using dimension reduction methods where the cells are represented in two or three dimensional space. Many tools are available to do this but visualising and cross-comparing these representations can be challenging, especially when cells are projected onto three dimensions which can be more informative for complex datasets. Here we present CellexalVR (www.cellexalvr.med.lu.se), a feature-rich, fully interactive virtual reality environment for the visualisation and analysis of single-cell experiments that allows researchers to intuitively and collaboratively gain an understanding of their data.

## Background

The analysis of single-cell RNAseq data (scRNAseq) is often performed using scripting, primarily using packages for the R/Python languages such as monocle [1], Seurat [2] and SCANPY [3] among others. A common step after pre-processing is dimension reduction (DR) where the cells are positioned in two/three dimensional space to visualise heterogeneity within the assayed populations. Many methods are available to do this (e.g, tSNE [4] and UMAP [5]), therefore several methods can be implemented during the course of an analysis and compared. Furthermore, each method has user-definable hyperparameters so outputs from the same methods also require comparison. Cells projected onto three dimensions can afford greater visual power to resolve the spatial arrangement of the cells and the clusters they form [6], particularly when cell-types are similar but distinct.

Many tools are available to visualise scRNAseq data [7], and published single-cell datasets are often accompanied by a webtool, where 3D DR plots can be explored in a rudimentary fashion [8,9]. The visualisation of 3D projected data is currently restricted to conventional computer monitors which have several shortcomings. The plots often have limited viewing angles due to them having only a single point of rotation at the centre, making it difficult to view populations on the periphery. Importantly, only one projection can be loaded in the same window meaning direct comparisons between several 3D reductions cannot be made easily. Another drawback is often these plots are not interactive, so for example, selecting cells for further analysis is not possible.

Here we present CellexalVR, a virtual reality (VR) platform that overcomes these barriers. VR is emerging as a potent tool to visualise scientific data, and its application thus far has been focused on image/volumetric data [10–12]. By placing all DR plots and associated metadata in VR we have created an immersive and collaborative environment to explore and analyse scRNAseq experiments.

## Results

As CellexalVR is based in virtual reality, it provides the ultimate environment to explore scRNAseq data, particularly when it has been arranged in three-dimensional space. To demonstrate the advantage of embedding cells into three dimensions versus two, Figure 1 (a) shows two subtypes of mesoderm overlapping during mouse gastrulation [13] in a UMAP using two-dimensions, however, projecting onto three-dimensions (b) shows them to be transcriptionally distinct. We tested this systematically by comparing the overlap between cell-types and clusters pairwise using entropy as a measure of mixing, where for (c) celltypes and (d) clusters, reduction to three-dimensions resulted in less overlap than in two-dimensions. This observation applied when using both UMAP (lower half) and tSNE (upper). We saw this trend consistently when applying this procedure to two further large datasets where we also used convex hulls as a second measure of overlap (Figure S1 and S2).

**Figure 1:**
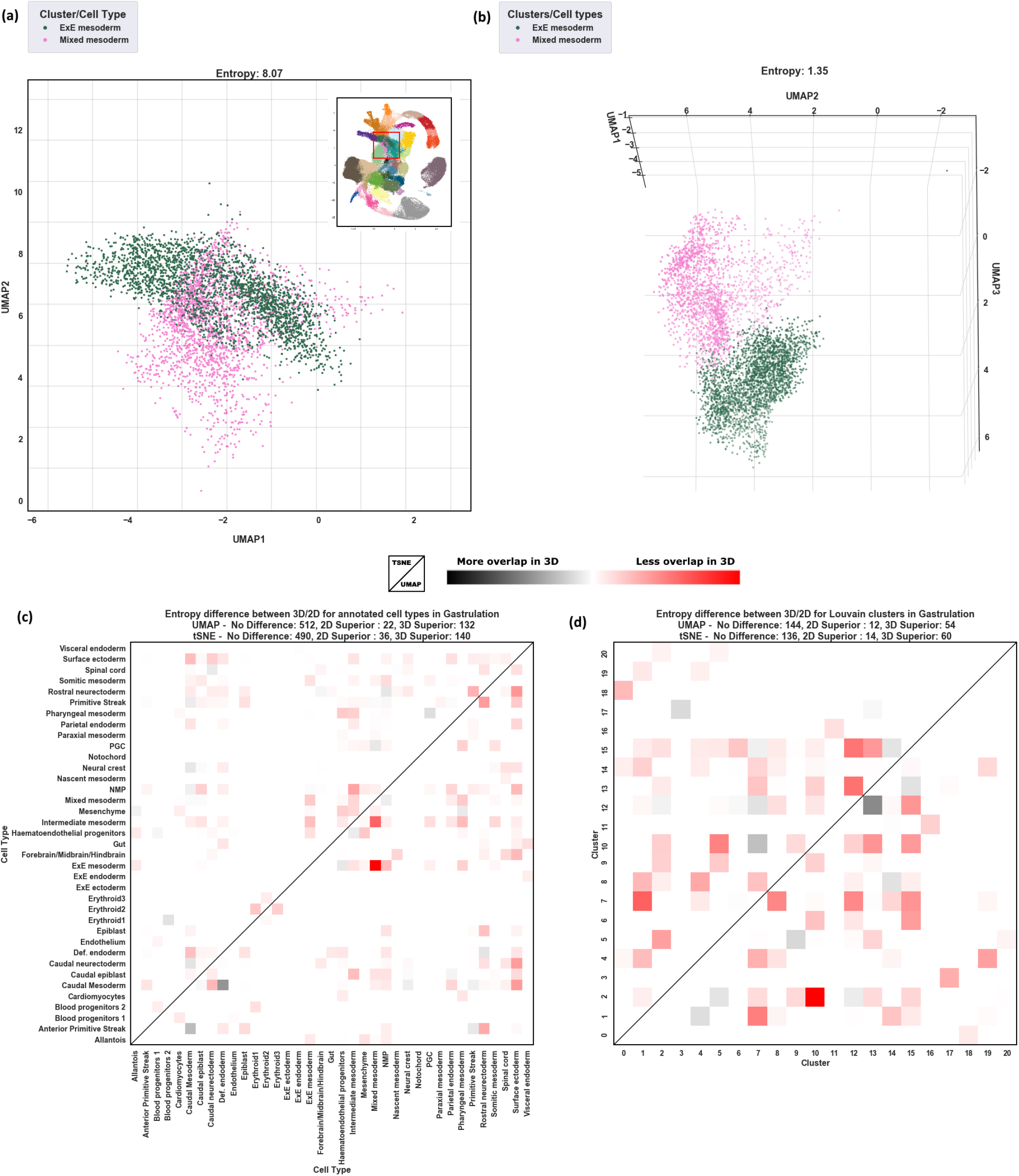
**(a)** Two mesoderm types from [13] overlap in a UMAP when projected onto two dimensions, where, in **(b)** they are transcriptionally distinct when projected onto three. **(c)** shows a pairwise comparison of cell type overlap using tSNE (upper diagonal) and UMAP (lower diagonal). Colour shows the difference in entropy where red signifies lower cell type collision/entropy in the 3D projection and grey denotes lower cell type collision in the 2D projection, where, it can be seen projection onto 3D overall has greater resolving power over 2D. **(d)** Shows the same pairwise test of overlap using a Louvain clustering of the cells.

In CellexalVR, DR plots can be interacted with intuitively, for example by grasping and moving them to gain any view required as if they were a physical object being held in one’s hand. Multiple DR plots can be visualised simultaneously and cross-compared with ease, for example, cells of interest in a tSNE plot can be captured with a hand gesture and traced to their counterparts in a UMAP allowing the user to directly visualise the differences between the two reduction methods.

At a minimum CellexalVR should be provided with a gene expression matrix and at least one set of DR coordinates (2D or 3D). CellexalVR will also import cell surface marker intensities captured during index sorting/CITEseq and categorical metadata for cells and genes. Figure 2b-f shows a selection of features currently available. DR plots can be coloured according to gene expression selected using a keyboard in the virtual environment, here the expression of *Gja1* in a tSNE and UMAP of 37k cells from aging mouse brain [14] (Figure 2b). Cells can be freehand captured into groups by passing them through a selection tool attached to the controller where they are coloured as they pass through. (Figure 2c). Selecting a new colour using the touchpad initiates a new group. Once two or more groups have been defined the user can, for example, calculate and display a heatmap of differentially expressed genes (Figure 2f left), or generate networks of transcription factors that can be cross compared (Fig S1). Cells captured in one DR plot can be traced to their counterparts in others to visually assess the differences (Figure 2d). Here, cells captured from a DDRTree of mouse hematopoietic stem and progenitor cells [9] split into two groups in a tSNE made from the same cells. Furthermore, using the selection tool the cells forming the split can be captured using the centrally generated graph to then determine which genes are differentially expressed between the two. Cells can also be coloured by user-provided attributes and for dense maps they can be rendered to a separate plot embedded within a skeleton of the data to better reveal their position within the total graph (Figure 2e). Here, two classes of mesoderm from 116k cells assayed during mouse gastrulation [13] are shown where the interface between these two cell-types is much clearer to see when spawned separately.

**Figure 2:**
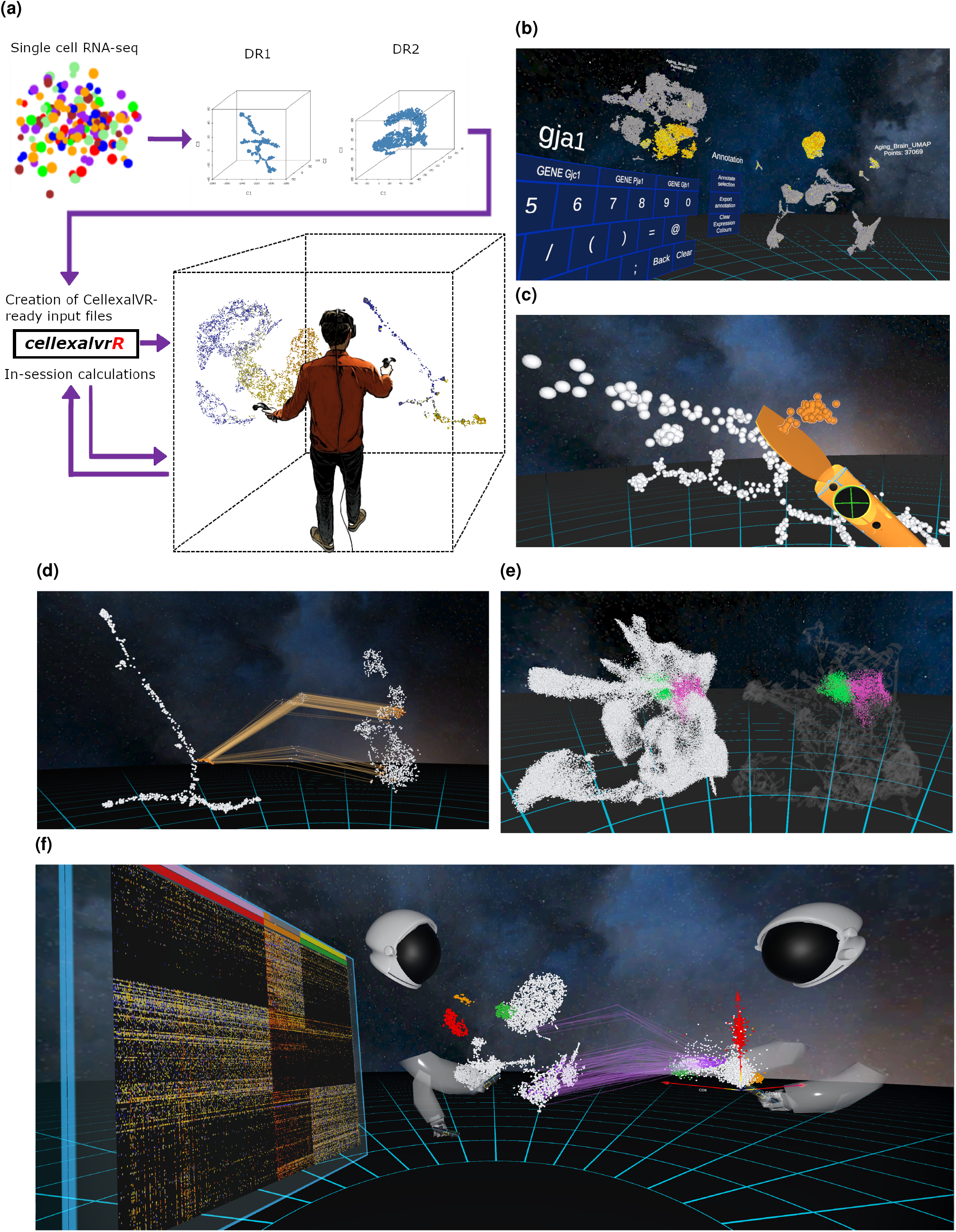
**(a)** Schematic overview of CellexalVR. Single-cell data is processed using user-preferred pipelines/methods and then exported using the cellexalvrR package to produce the files required by CellexalVR. **(b)** A tSNE and UMAP of cells from the aging brain coloured by the expression of Gja1. **(c)** Cells can be freehand captured by passing them through the selection tool and clicking the touchpad changes colour/initiates a new group. **(d)** Cells selected from the DDRTree plot (left) are tracked to their counterpart cells in the tSNE plot (right), generating a new graph of points between the two that can also be selected on. **(e)** Cell types can be coloured and spawned in a separate graph if they are difficult to see in dense DR plots. Here, two-types of mesoderm are rendered within a skeleton of the data (right). **(f)** Multiuser mode showing two users in the same CellexalVR session analysing the same data together.

CellexalVR has been built using video game technology, and we have leveraged this to develop multiuser mode where several individuals can meet in the same CellexalVR session and analyse the same data together, a big advantage given many datasets are generated collaboratively over different geographic sites. Figure 2f shows two users analysing data from cord blood mononuclear cells from a CITEseq experiment [15] where the right-most user has chosen three surface markers to plot against each other, and has tracked a selection of these cells to the tSNE of the RNA data. The left user has selected groups from the UMAP and created a heatmap of differentials. Each participant will need access to a VR headset, controllers, workstation, and a copy of the input data files. It is also possible for users to join a session using just a desktop computer by navigating the session using a keyboard and mouse. These participants cannot be seen by the VR active users or interact with the session objects, hence we call this “spectator mode”. All voice is transmitted using a third-party application of the user’s choice (Skype/Discord/Zoom to name a few).

We have also developed a novel way to visualise output from RNA velocity [16] when the user has calculated trajectories from a 3D projection of the cells. The velocity of each cell is shown as an animated process where an arrow/point travels the length of the calculated trajectory. The arrows are spawned randomly to avoid crowding, and the overall effect shows a “flow” of cells as they traverse the dimension reduced data. This removes any need to average velocities for plotting on a 2D map where subtleties can be lost depending on the grid size chosen. These trajectories can also be coloured by cell attribute/gene expression adding a further dimension to the information that can be simultaneously visualised. This feature is demonstrated using 142K cells from human fetal liver [17] where we show the trajectories being coloured by gene expression level and cluster membership (https://youtu.be/iFpqRLpCRms).

User comfort has been a key priority during the development of CellexalVR which has been designed to eliminate VR induced motion sickness which is a known issue. The default environment is relatively dark which is easy on the eyes and provides high contrast against the cells. Other environments are available from the configuration menu which also allows the setup a custom colour palette, and change the size and rendering quality of the cells. A dedicated VR headset and controllers provide a smooth, high resolution view of the environment with a wide field of view which is fully tactile and fast. Importantly, CellexalVR is comfortable to use for extended periods of time (1 hour +) which is essential for the thorough analysis of complex data. CellexalVR can also export heatmaps, gene networks, cell annotations, screenshots (via the camera tool), and results generated during a session leading to productive and documented sessions. For those not familiar with VR we have created an interactive training session where the user is familiarised with a VR environment and shown how to perform basic functions in CellexalVR.

## Preparing data for CellexalVR

CellexalVR requires several input files which are detailed on the project website, but to make it easy we have created an R package (cellexalvrR) to help prepare them. Those using Seurat can use our “as_cellexalvrR” function which will produce a cellexalvrR object that is exported using “exprort2cellexalvr”. Those using Scanpy can import and convert their AnnData object to a Seurat object (using seurat-disk) and export from there. To enable RNA velocity visualisation one supplies coordinates denoting the destination of each cell alongside the coordintes of the DR method used. All details and tutorials regarding use of the R package can be found at https://cellexalvr.med.lu.se/cellexalvrr-vignette. Importing data from multiple analysis platforms is a key challenge and one that we will continue to work on.

## Data types handled by CellexalVR

The data supplied to CellexalVR is flexible regarding the source. The expression matrix and DR coordinates used as input can be derived from other experiment types. For example, gene activities from single-cell ATACseq (Figure S4), and we have also used CellexalVR to visualise CyTOF data. In the near future we will routinely see multi-modal data from the same cell (such as RNA and epigenetic information), and CellexalVR will be the perfect environment to unify all visualisations from these complex and information rich experiments. There is also a clear application of CellexalVR to spatial transcriptomics experiments which will form a future update.

## Comparison to other software

There are currently two other projects in preprint that allow users to visualise single-cell data in VR, starmapvr [18] and singlecellvr [19], that between them cover a range of features.

The main difference between CellexalVR and these projects is the underlying technology being used. Both starmapvr and singlecellvr utilise Google cardboard where a mobile phone is placed inside a visor to form a rudimentary headset. These systems have three degrees of freedom (3-DoF) tracking which means only head rotation is tracked. As *position* is not tracked, a keyboard (or voice control in the case of starmapvr) is needed to walk around the scene. Also, the absence of tracked and visible hand controllers means object interaction has to be done by other means (keyboard, voice). CellexalVR uses a dedicated VR system connected to a workstation. Head rotation *and* position is tracked (6-DoF), and with tracked controllers it places the users hands in the same environment for effortless object manipulation (grabbing/moving/rotation). Another key difference is image quality. Cardboard systems render the scene on the phone, whereas a dedicated VR system renders the scene on workstation/laptop with a more powerful GPU, therefore will show far more detail with less latency.

starmapvr and singelcellvr represent a good attempt at making 3D visualisation more accessible, however, using the capabilities of a true VR system means CellexalVR can provide a wider range of tools to pick apart single-cell data, and has scope for further expansion.

## Conclusions

The analysis and interpretation of scRNAseq experiments are heavily reliant on data visualisation, and we have shown that virtual reality is the perfect medium in which to fully scrutinise this complex data in a manner which is intuitive and fully collaborative. VR technology will continue to develop, as will CellexalVR as we improve and introduce functionality to keep apace of VR and single-cell technology.

## Methods

CellexalVR comprises of two main components. First is a VR environment which has been implemented using the Unity game engine (https://unity.com/). CellexalVR uses several Unity assets that are libraries containing scripts that handle parts of the program logic. SteamVR (https://steamcommunity.com/steamvr) and OpenVR (https://github.com/ValveSoftware/openvr) handle communication between the computer, headset and controllers, and VRTK (https://github.com/thestonefox/VRTK) handles basic interaction logic such as the grabbing of objects. In order to accommodate a large number of points/cells a new collision detection system was engineered (supplementary methods).

The second component is an R package called cellexalvrR that 1) provides simple to use functions to export scRNAseq data from an R session to a set of files CellexalVR can import, and 2) performs back-end calculations during a CellexalVR session (differential expression and correlation analysis among others). Compartmentalising CellexalVR this way means bioinformaticians can alter the R package to customise the underlying methods without needing knowledge of C# (the language used by Unity).

Multi-user mode is facilitated via the Photon Unity Networking (PUN, https://www.photonengine.com/en-US/PUN) that works by sending packages that contain information about events between the users through Remote Procedure Calls (RPCs). The RPCs ensure that each user’s session is synchronised with all others. The data files are *not* transmitted over RPCs, therefore each user must have a copy of the data being analysed locally on the workstation they are using. Head models were downloaded from NASA (https://nasa3d.arc.nasa.gov) and the arm models were made in Blender (https://www.blender.org/).

Overlaps between cell-types and clusters were done using two methods, 1) using the Shannon Index and, 2) defining the groups as convex hulls. Details are in the supplementary methods with links to the source code.

## Availability and requirements

CellexalVR is available from the project website at https://cellexalvr.med.lu.se along with extensive documentation and video tutorials. The data used here is also available from the project website and cited references.

The source code for the CellexalVR is available at https://github.com/sonejilab/cellexalvr which is coded in C#, and the accompanying R package cellexalvrR can be installed directly from GitHub https://github.com/sonejilab/cellexalvrR. For those wanting to modify CellexalVR, we have a developers guide at https://cellexalvr.med.lu.se/programmers-guide.

### Hardware

Users require a VR ready workstation/laptop with a suitable graphics card (Steam recommends a GTX1060 or higher) running Windows 10. Regarding VR equipment, CellexalVR was developed on the HTC Vive/Vive Pro and Valve Index headsets, and the documentation we provide describes how to operate the HTC Vive controllers. Documentation for others will follow, but CellexalVR has been tested successfully with the Valve Index controllers too. These VR systems are readily available, and are priced for the home consumer.

## Authors’ contributions

O.L and J.R developed CellexalVR, created the project website, wrote the online documentation and, prepared figures. J.P developed CellexalVR. P.D contributed code and technical assistance. S.L and S.S created cellexalvrR and M.W provided technical assistance. S.S conceived the study. O.L, J.R and S.S wrote the paper. O.L and J.R contributed equally.

## Acknowledgements

We thank Rasmus Olofzon, Kristian Berg, Daniel Hellström, Daniel Cheveyo, Arvid Carlman, and Christopher Nilsson from the LTH, Lund University for their work on the prototype. Steve Taylor at the CBRG, Oxford University, Ivan Imaz-Rosshandler at the Cambridge Stem Cell Institute and members of the Lund Stem Cell Centre for testing and feedback. We also thank the good people of Stack Overflow. O.L and J.R are funded by the Knut and Alice Wallenberg Foundation (KAW2014.0098) and Cancerfonden (CAN2018-565). S.L and S.S are funded by StemTherapy which is funded by the Swedish Government.

