## Supplemental figures and methods for "CellexalVR: A virtual reality platform to visualise and analyse single-cell data"

### Supplementary Figures

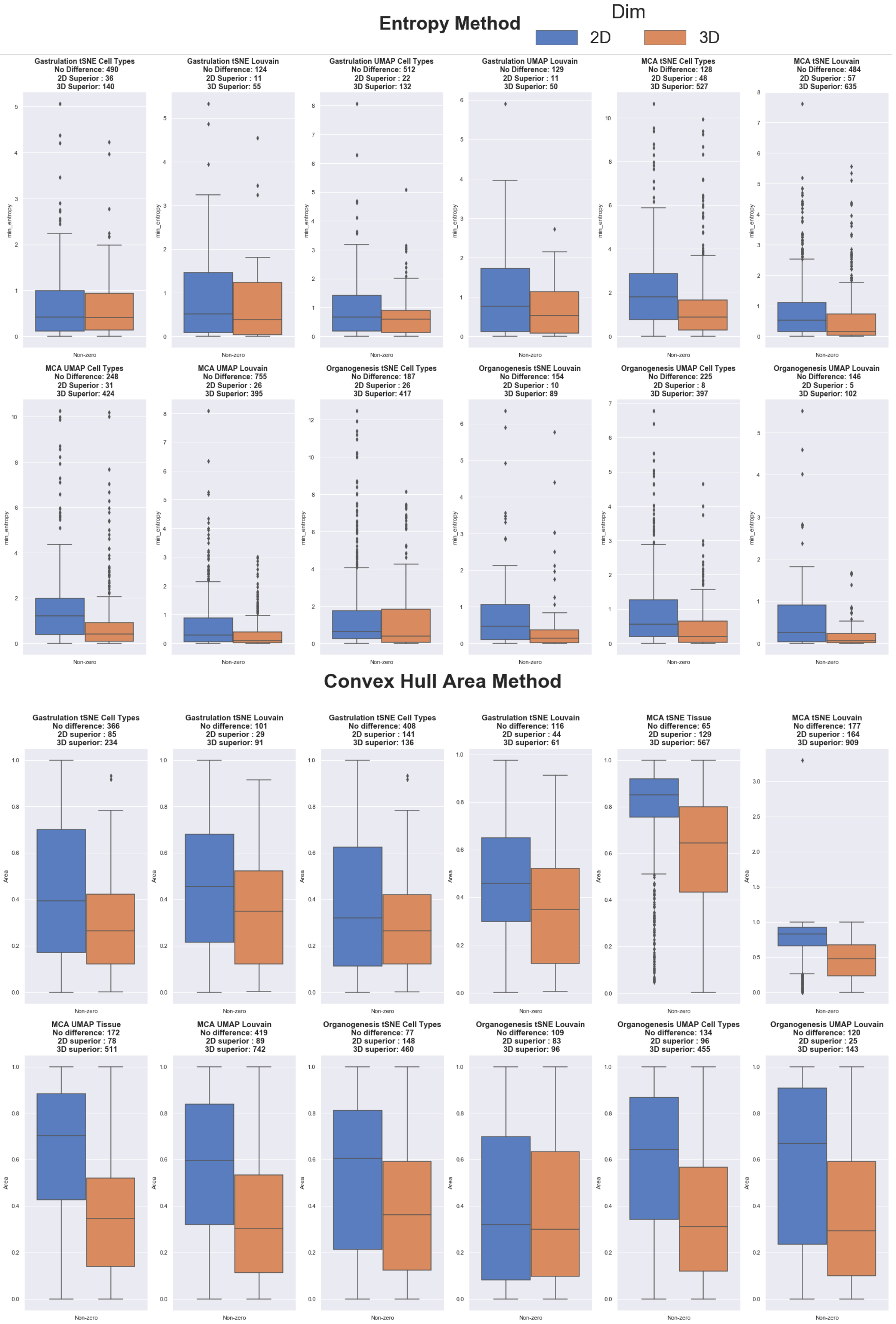


Figure S1: A comparison of 2D (blue boxes) vs 3D (orange boxes) dimension reduction of scRNAseq using three datasets implementing UMAP and tSNE. Boxplots show the distribution of entropies (top 12), or, area overlap of convex hulls (bottom 12) which were calculated comparing pairwise the overlap between cell types and between Louvain clusters. Overall we observe projecting the cells onto 3 dimensional data will allow close but distinct populations to be visually resolved to a greater extent when compared to 2D projections.

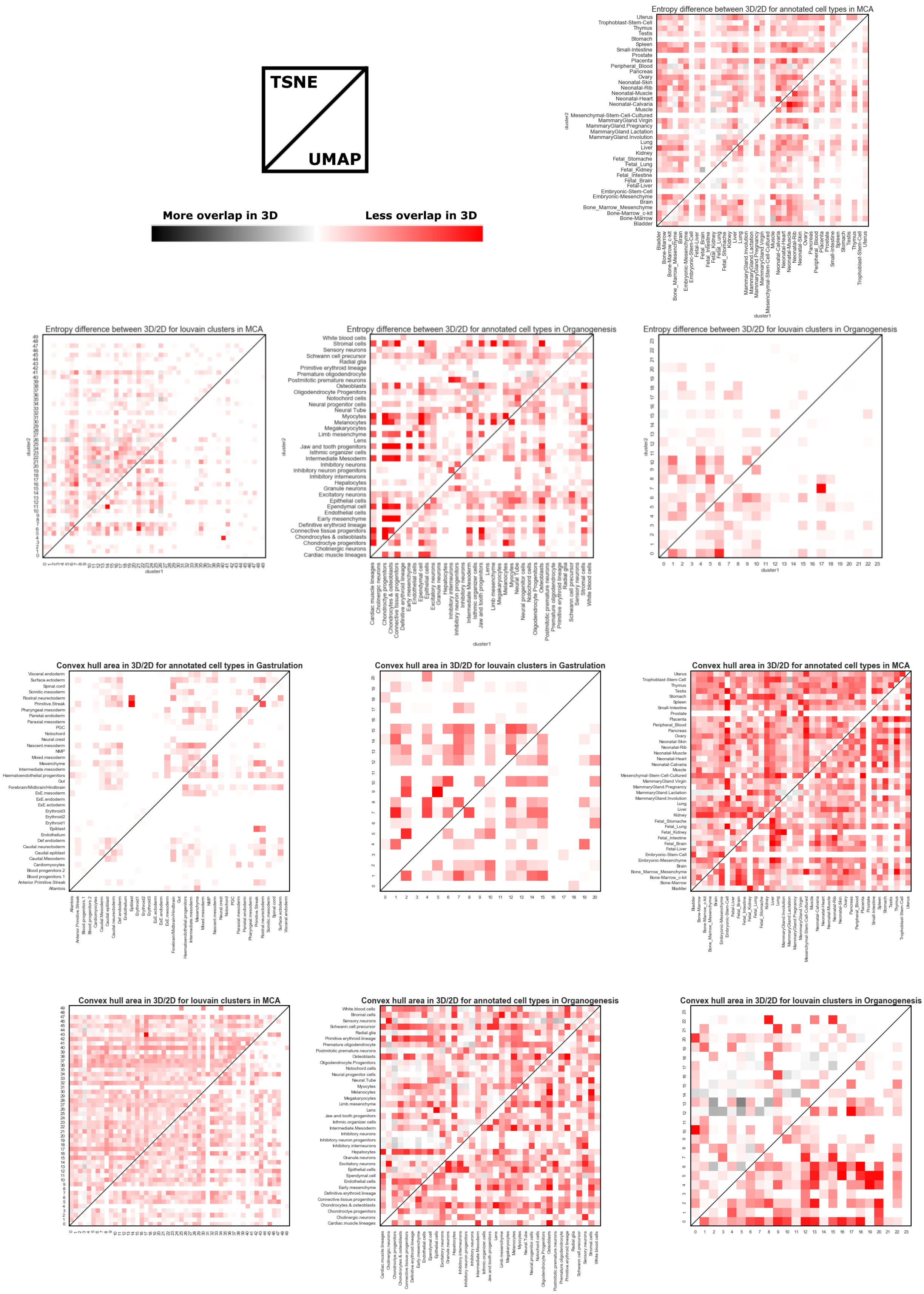


Figure S2: Heatmaps of the data shown in FigS1. Each shows the pairwise difference in entropy (top four), or, area overlap of convex hulls (bottom 6) which were calculated comparing the overlap between cell types and between Louvain clusters. Red denotes less cell mixing in 3D reduced data which is the dominant trend within these datasets regardless of DR method used.

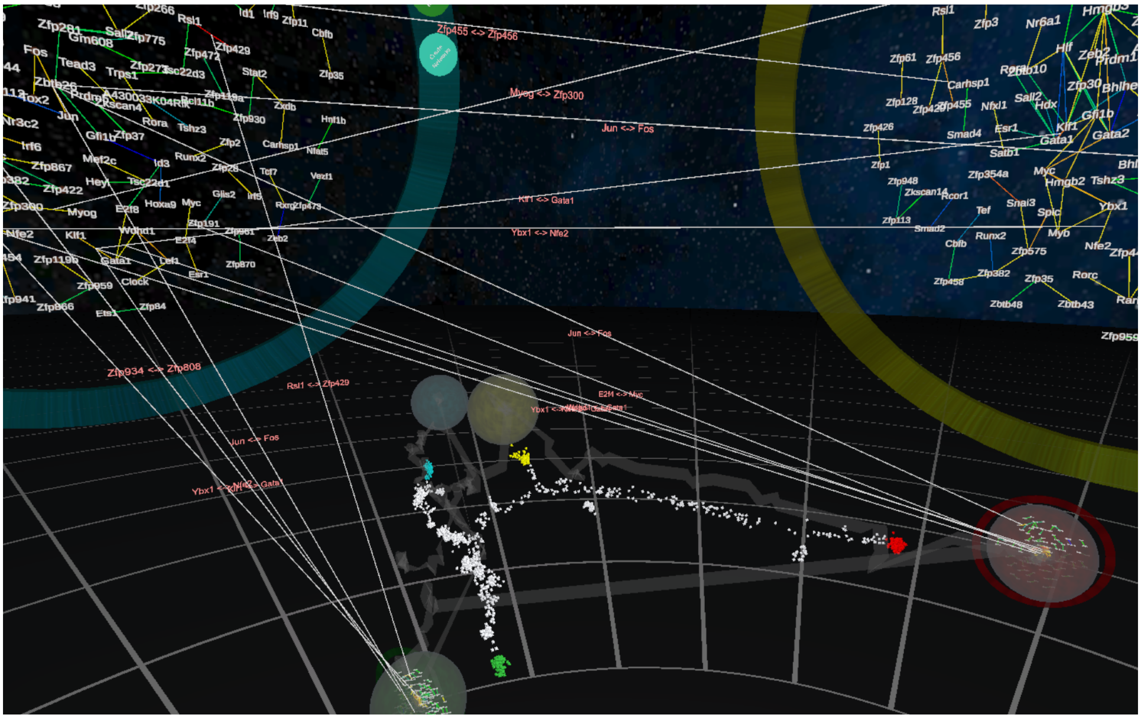

Figure S3: TF-TF correlation networks that have been calculated for each of the groups captured in the dimension reduced plot in the background. Each network is rendered in a sphere that is embedded in a abstract representation of the DR graph to show each networks originating group. Here, two networks have been clicked and expanded, and common TF-TF pairs have been toggled and highlighted to show where commonalities lie between groups

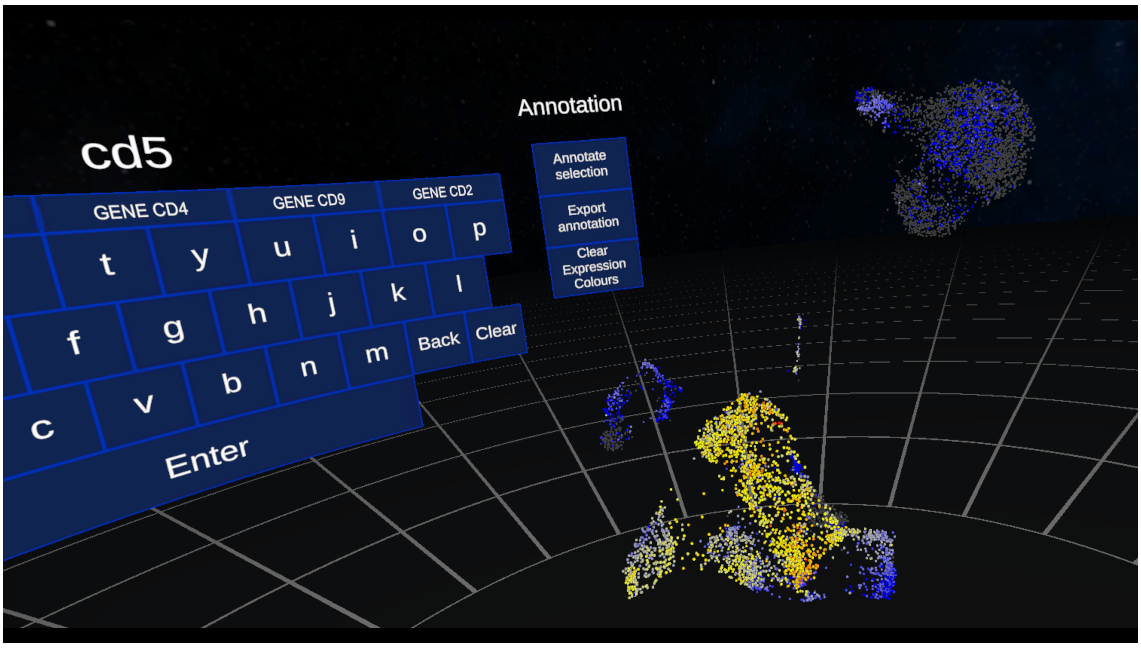


Figure S4: A UMAP generated from single-cell ATACseq from 7866 PBMC cells. Here, each cell is coloured by the activity of cd5. Data was processed according to the Seurat documentation `https://satijalab.org/seurat/v3.2/atacseq_integration_vignette.html` and exported using cellexalvrR.

**Movie M1**: RNA Velocity in CellexalVR. Reads from the Liver Cell Atlas were aligned using kallisto-bustools [1] using loompy's "fromfq" function (http://loompy.org/). The resulting loom file was processed through scvelo [2] to calculate the RNA velocities and the output files were made in R using our cellexalvrR package. Trajectories were based on two 3D embeddings, PHATE [3] and UMAP. Each cell emits an arrow that travels the length of the calculated vector. The the longer the trajectory the redder the arrow. Arrows can be changed to small particles for easier viewing on dense plots, and both emission types can be coloured by gene expression and cell attribute.

**Supplementary Methods**
CellexalVR is built using Unity (https://unity.com/) that handles tasks such as rendering the frames that are displayed on the computer's monitor and in the user's headset, collecting input from the keyboard and the controllers and forwarding events that trigger certain actions and handling all physics simulation. Unity comes with an editor which is the primary development environment CellexalVR was created with.

In order to accommodate a high-number of cells while retaining the stipulation that each cell must be individually selectable, we wrote a new collision detection system utilising Octrees [4] and raycasts. This is described at https://www.cellexalvr.med.lu.se/programmers-guide along with a detailed overview of the code structure and how to navigate it for those wishing to modify it. The code is hosted on GitHub at https://github.com/sonejilab/cellexalvr.

Input files for CellexalVR are produced in R using our cellexalvrR package (https://github.com/sonejilab/cellexalvrR). A guide is provided at https://www.cellexalvr.med.lu.se/cellexalvrr-vignette where we give working examples showing how to make a CellexalVR object and export it to files that CellexalVR will import. We also give an example of how to make and export RNA velocity embedding in 3D for visualisation. A detailed description of the file formats can be seen at https://cellexalvr.med.lu.se/manual_introduction. During export the following files are produced:

- **An R object containing all data**. This is loaded into an R server session when the data is loaded in CellexalVR in preparation for in-session calculations.
- **An SQLite database** that Unity uses to draw expression values from when colouring cells by gene expression and rendering heatmaps. SQLite is used because it carries significant speed advantages.
- **Basic text files** that are used to populate the environment (DR plots and metadata) when data is loaded.

In-session calculations are also performed in R using cellexalvrR. When a data set is loaded an R server is created and initialised with the expression data. Differentially expressed genes for heatmaps are identified using the Wilcox test (implemented in C++ for speed) and clustering is performed using hierarchical clustering for the top $N$ genes (default is 250). Transcription factor (TF) networks within selected groups of cells can be calculated using the rho value (default) from the propr package, or partial correlations from the ppcor package. The top 130 interactions are returned by default, but is user configurable. All heatmaps and networks are rendered in the CellexalVR UI. TFs are defined as those in the AnimalTFDB database (http://bioinfo.life.hust.edu.cn/AnimalTFDB/).

**Data formats**. CellexalVR requires correctly formatted input files. This process is greatly simplified by using the R package cellexalvrR that will generate the required files, R object, and SQLite database of expression data. At a minimum CellexalVR should be provided with:

- ***A matrix of gene expression data (C cells x G genes)***. This is processed by cellexalvrR to an SQLite database that CellexalVR queries when needed.
- ***At least one set of DR coordinates placing the cells in 3D space (C cells x 3)***. This can be from any DR methods the user deems suitable and CellexalVR will accept more than one DR table. These are exported as 4-column text files (*.mds) with the Cell ID in the first column. If RNA Velocity trajectories have been added these will be 7-column .mds files.

In addition to these, further optional files can be imported. These are:

- ***Surface marker intensities (C cells x S surface markers)***. These are recorded when cells have been index sorted. If using CITEseq, these expression values go into this table.
- ***Cell type information (C cells x T types)***. These allow the user to label each cell as being of a certain type, which can then be displayed in the CellexalVR session. Cells are marked as belonging to a class with a "1", or "0" otherwise.
- ***Metadata for cells (C cells x M meta)***. Further labels for cells, for example cell-cycle stage.
- ***Metadata for genes (G genes x M meta)***. For example, marking genes if they belong to a particular category such as transcription factors, epigenetic factors, or code surface proteins.

To export the necessary file from R, a cellexalvrR S4 object needs to be created first using the supplied functions. More details on how to export CellexalVR ready files can be seen at https://www.cellexalvr.

med.lu.se/cellexalvrr-vignette.

**Output from CellexalVR**. During a CellexalVR session the user has the option to document what they have done using various tools and methods:

- Capture images using the "Camera tool" button placed on the menu controller.
- Save text files of annotated Cell IDs when the user has used annotation mode.
- Images of heatmaps can be saved by clicking on the "Save" button to the left of the heatmap.
- Images of TF networks can be saved by clicking on the "Save as image" button on the surrounding border.
- An adjacency list of a network can be saved by clicking on the "Save as text file" button on the surrounding border.
- The modified R object is returned containing all the user captured groups.
- **An html report is generated** when the user clicks the "Save session" button on the menu, or when the program is exited. Everything the user has decided to save during a session is compiled into an HTML report with figures (heatmaps and networks) and associated p-values and FDR's for the differentially expressed genes. For gene lists, enriched Gene Ontologies are automatically calculated and tabulated using TopGO.

**Software requirements**. CellexalVR requires the following to work:

- R 3.6 or greater.
- Rtools (https://cran.r-project.org/bin/windows/Rtools/) which is used to compile sub-routines in cellexalvrR.
- Pandoc (https://pandoc.org/installing.html) which is used to compile HTML reports.
- cellexalvrR (https://github.com/sonejilab/cellexalvrR).
- SteamVR (which should be installed automatically when he headset is setup).

**Hardware**. CellexalVR was developed on a gaming class workstation comprising an Intel i7 processor, 16GB RAM, 1TB SSD, and an NVIDIA GTX1080 graphics card. We used the HTC Vive Pro and Valve Index headsets, and the documentation currently describes how to use the HTC Vive controllers, but CellexalVR will work with the Valve Index controllers too.

**2D/3D projection comparison**

Dimension reduction methods for data visualisation are prone to the introduction of distortions into data topology [5] and 3D projections of data have been shown to preserve the distances between the data-points better than their 2D counterparts [6]. Controlled user studies have shown that, compared to 2D, 3D projections improves the ability to discriminate datapoint clusters visually [7]. Furthermore, it has been shown that visual inspection of datapoints is better when 3D embeddings are explored interactively rather than as fixed axis animated movies [8].

Two different approaches were taken to measure the degree of overlap/separation between cell-types and clusters in 3D and 2D projections of scRNAseq data, 1) an entropy based method, and 2) measuring area/volume overlap between convex hulls. For example, two cell populations that seemingly overlap in a 2D projection can be resolved into separate populations when the cells are projected onto three dimensions instead, and this can be seen when rotating the 3D plot. To mimic this, for both the entropy and convex hull method we computationally rotate the 3D projections of the data and then "flatten" the cells to a 2D representation before calculating pairwise the overlap between cell type/clusters. We retain the value corresponding to the least overlap and compare that to the data when projected onto two dimensions only. In both entropy and hull methods we used UMAP and tSNE using three large and well annotated datasets; 116k cells from mouse gastrulation [9], 100k cells from mouse organogenesis [10], and, the Mouse Cell Atlas [11].

**Entropy Method** This approach uses the *Shannon Index* ($SI$) as the measure of overlap. To start, a grid is placed over the UMAP/tSNE dividing it into a number of different sectors. For each sector the $SI$ is calculated using the following:

$$SI = -\sum_{i=1}^{R} p_i \ln p_i \tag{1}$$

where $p_i$ is the proportion of cells belonging to the $i$th population, $R$ is the number of different populations, since we are doing pairwise comparisons $R = 2$. Lastly the $SI$ of each sector is summed together. To ensure the placement of the grid is not a major influential factor of the result, we shift the grid stepwise and use the position that gives the lowest overall $SI$ value. This is done for both the 2D and 3D setting.

As explained above, the 3D graphs are flattened to 2D after systematically rotating the graph to cover all angles. For each of these projection angles a grid is placed and the $SI$ is calculated the same way as in equation 1. The projection that gives the lowest $SI$ value is picked as the best viewing angle and this is the value that is then compared to the 2D graph. If the value is lower for the best viewing angle in the 3D graph it means the third dimension helped separate these populations since you were able to find an angle where the cell populations or clusters had less overlap compared to the 2D projection. Code for this method can be obtained at https://github.com/sonejilab/entropy_2dvs3d

**Convex Hull Area Method** In a 2D projection of the data, convex hulls were calculated for each defined cell type or cluster. The convex hulls were then compared pairwise and the overlap was calculated in terms of area. The area of the overlap between the two convex hulls was calculated as a percentage of the smallest hull in terms of area. In 3D projections, the populations were first flattened onto 2 dimensions through 2048 different angles in order to emulate what a user sees from a given angle. For each pair of cell type/clusters the viewing angle with the least amount of overlap in terms of area was found using an exhaustive search. A convex hull of the overlap was calculated and populations were compared pairwise as for the 2D setting.

The convex hulls were calculated using the Quickhull algorithm [12] and the overlapping area between two given convex hulls were calculated [13]. Code for this method can be obtained from https://github.com/sonejilab/2d_vs_3d.

### References


[1] Páll Melsted, A. Sina Booeshaghi, Fan Gao, Eduardo Beltrame, Lambda Lu, Kristján Eldjárn Hjorleifsson, Jase Gehring, and Lior Pachter. Modular and efficient pre-processing of single-cell RNA-seq. *bioRxiv*, 2019.

[2] Volker Bergen, Marius Lange, Stefan Peidli, F. Alexander Wolf, and Fabian J. Theis. Generalizing RNA velocity to transient cell states through dynamical modeling. *bioRxiv*, 2019.

[3] Kevin R. Moon, David van Dijk, Zheng Wang, William Chen, Matthew J. Hirn, Ronald R. Coifman, Natalia B. Ivanova, Guy Wolf, and Smita Krishnaswamy. PHATE: A Dimensionality Reduction Method for Visualizing Trajectory Structures in High-Dimensional Biological Data. *bioRxiv*, 2017.

[4] Donald Meagher. Geometric modeling using octree encoding. *Computer Graphics and Image Processing*, 1982.

[5] Benoit Colange, Laurent Vuillon, Sylvain Lespinats, and Denys Dutykh. Interpreting Distortions in Dimensionality Reduction by Superimposing Neighbourhood Graphs. In *2019 IEEE Visualization Conference, VIS 2019*, 2019.

[6] Danilo B. Coimbra, Rafael M. Martins, Tácito T.A.T. Neves, Alexandru C. Telea, and Fernando V. Paulovich. Explaining three-dimensional dimensionality reduction plots. *Information Visualization*, 2016.

[7] J. Poco, R. Etemadpour, F. V. Paulovich, T. V. Long, P. Rosenthal, M. C.F. Oliveira, L. Linsen, and R. Minghim. A framework for exploring multidimensional data with 3D projections. *Computer Graphics Forum*, 2011.

[8] Harald Sanftmann and Daniel Weiskopf. 3D scatterplot navigation. *IEEE Transactions on Visualization and Computer Graphics*, 2012.

[9] Blanca Pijuan-Sala, Jonathan A Griffiths, Carolina Guibentif, Tom W Hiscock, Wajid Jawaid, Fernando J Calero-Nieto, Carla Mulas, Ximena Ibarra-Soria, Richard C V Tyser, Debbie Lee Lian Ho, Wolf Reik, Shankar Srinivas, Benjamin D Simons, Jennifer Nichols, John C Marioni, and Berthold Göttgens. A single-cell molecular map of mouse gastrulation and early organogenesis. *Nature*, 2019.

[10] Junyue Cao, Malte Spielmann, Xiaojie Qiu, Xingfan Huang, Daniel M. Ibrahim, Andrew J. Hill, Fan Zhang, Stefan Mundlos, Lena Christiansen, Frank J. Steemers, Cole Trapnell, and Jay Shendure. The single-cell transcriptional landscape of mammalian organogenesis. *Nature*, 2019.

[11] Xiaoping Han, Renying Wang, Yincong Zhou, Lijiang Fei, Huiyu Sun, Shujing Lai, Assieh Saadatpour, Zimin Zhou, Haide Chen, Fang Ye, Daosheng Huang, Yang Xu, Wentao Huang, Mengmeng Jiang, Xinyi Jiang, Jie Mao, Yao Chen, Chenyu Lu, Jin Xie, Qun Fang, Yibin Wang, Rui Yue, Tiefeng Li, He Huang, Stuart H. Orkin, Guo Cheng Yuan, Ming Chen, and Guoji Guo. Mapping the Mouse Cell Atlas by Microwell-Seq. *Cell*, 2018.

[12] C. Bradford Barber, David P. Dobkin, and Hannu Huhdanpaa. The Quickhull Algorithm for Convex Hulls. *ACM Transactions on Mathematical Software*, 1996.

[13] Joseph O'Rourke, Chi Bin Chien, Thomas Olson, and David Naddor. A new linear algorithm for intersecting convex polygons. *Computer Graphics and Image Processing*, 1982.
